# Network analysis of ten thousand genomes shed light on *Pseudomonas* diversity and classification

**DOI:** 10.1101/2021.08.16.456539

**Authors:** Hemanoel Passarelli-Araujo, Glória Regina Franco, Thiago M. Venancio

**Affiliations:** Departamento de Bioquímica e Imunologia, Instituto de Ciências Biológicas, Universidade Federal de Minas Gerais, Belo Horizonte, MG, Brazil; Laboratório de Química e Função de Proteínas e Peptídeos, Centro de Biociências e Biotecnologia, Universidade Estadual do Norte Fluminense Darcy Ribeiro, Campos dos Goytacazes, RJ, Brazil

**Keywords:** Phylogenomics, Pseudomonads, Taxonomy, Community detection

## Abstract

The growth of sequenced bacterial genomes has revolutionized the assessment of microbial diversity. *Pseudomonas* is a widely diverse genus, containing more than 254 species. Although type strains have been employed to estimate *Pseudomonas* diversity, they represent a small fraction of the genomic diversity at a genus level. We used 10,035 available *Pseudomonas* genomes, including 210 type strains, to build a genomic distance network to estimate the number of species through community identification. We identified taxonomic inconsistencies with several type strains and found that 25.65% of the *Pseudomonas* genomes deposited on Genbank are misclassified. The phylogenetic tree using single-copy genes from representative genomes in each species cluster in the distance network revealed at least 14 *Pseudomonas* groups, including *P. alcaligenes* group proposed here. We show that *Pseudomonas* is likely an admixture of different genera and should be further divided. This study provides an overview of *Pseudomonas* diversity from a network and phylogenomic perspective that may help reduce the propagation of mislabeled *Pseudomonas* genomes.

## INTRODUCTION

Biological networks have been an essential analytical tool to better understand microbial diversity and ecology^1, 2^. A network is a set of connected objects, in which objects can be represented as nodes and connections as edges. Networks provide a simple and powerful abstraction to evaluate the importance of individual or clustered nodes in maintaining a given system. Coupled with whole-genome sequencing, it can refine our knowledge about genetic relationships of diverse bacteria such as *Pseudomonas*.

*Pseudomonas* is a genus within the *Gammaproteobacteria* class, whose members colonize aquatic and terrestrial habitats. These bacteria are involved in plant and human diseases, as well as in biotechnological applications such as plant growth-promotion and bioremediation^3^. The genus *Pseudomonas* was described at the end of the nineteenth century based on morphology, and its remarkable nutritional versatility was recognized thereafter^4^. The metabolic diversity of pseudomonads, combined with biochemical tests to describe species, culminated in a chaotic taxonomic situation^4^.

In 1984, the genus was revised and subdivided into five groups based on DNA-DNA and rRNA-DNA hybridization^5^, with group I retaining the name *Pseudomonas*. Over the past 30 years, other molecular markers such as housekeeping genes have been used to mitigate the issues of *Pseudomonas* taxonomy^6, 7, 8^. Based on the 16S rRNA gene sequences, the genus is divided into three main lineages represented by *Pseudomonas pertucinogena*, *Pseudomonas aeruginosa*, and *Pseudomonas fluorescens*^9^. These lineages comprise groups of different species – both lineages and groups receive the name of the representative species. Currently, there are 254 *Pseudomonas* species with validated names according to the List of Prokaryotic Names with Standing in the Nomenclature (LPSN)^10^. However, although the genus division into lineages and groups has facilitated the classification of new species, the remnants of the *Pseudomonas* misclassification still linger in public databases^11, 12^.

The explosion in the availability of complete genomes for both cultured and uncultured microorganisms has improved the classification of several bacteria, including *Pseudomonas*^8, 13^. One of the gold standards for species circumscription is the digital whole-genome comparison by Average Nucleotide Identity (ANI)^14^. Since using only genomes from type strains might bias and provide an unrealistic picture of microbial diversity, we aimed to estimate the *Pseudomonas* diversity using all available genomes through a network approach. Here, we provide new perspectives on *Pseudomonas* diversity by exploring the topology of the genomic distance network and the phylogenetic tree from representative genomes. This work also provides novel insights into the misclassification and phylogenetic borders of *Pseudomonas*.

## RESULTS

### Dataset collection

We obtained 11,025 genomes from GenBank in June 2020. After evaluating the quality of each genome (see methods for more details) and removing fragmented genomes, 10,035 genomes passed in the 80% quality threshold (Figure S1). The size of the retrieved genomes ranged from 3.0 to 9.4 Mb. We used 238 type strains with available genomes and names validly published according to the *List of Prokaryotic Names with Standing in Nomenclature* in March 2021. The genome size and GC content of type strains ranged from 3,022,325 bp and 48.26% (*P. caeni*) to 7,375,852 bp and 62.79% (*P. saponiphila*) (Table S1). According to the NCBI classification, the top four abundant species in our dataset are *P. aeruginosa* (n = 5,088), *P. viridiflava* (n = 1,509), *Pseudomonas sp.* (n = 1,083), and *P. syringae* (n = 435) (Table S2).

### Genome-based analysis reveals the presence of synonymous *Pseudomonas* species

The misclassification of some *Pseudomonas* type strains has been reported by several studies^8, 15, 16, 17^. Type strains play an essential role in taxonomy by anchoring species names as unambiguous points of reference^18^. In this context, the term “synonym” refers to the situation where the same taxon receives different scientific names. We used 238 type strain genomes to evaluate the presence of synonymous species in *Pseudomonas.* The ANI was computed for all type strains to construct an identity network further used to check the linkage between genomes based on a 95% ANI threshold (Figure 1). Since 95% has been accepted as species delimitation threshold^14^, connections between type strains indicate synonymous names or subspecies.

**Figure 1.**
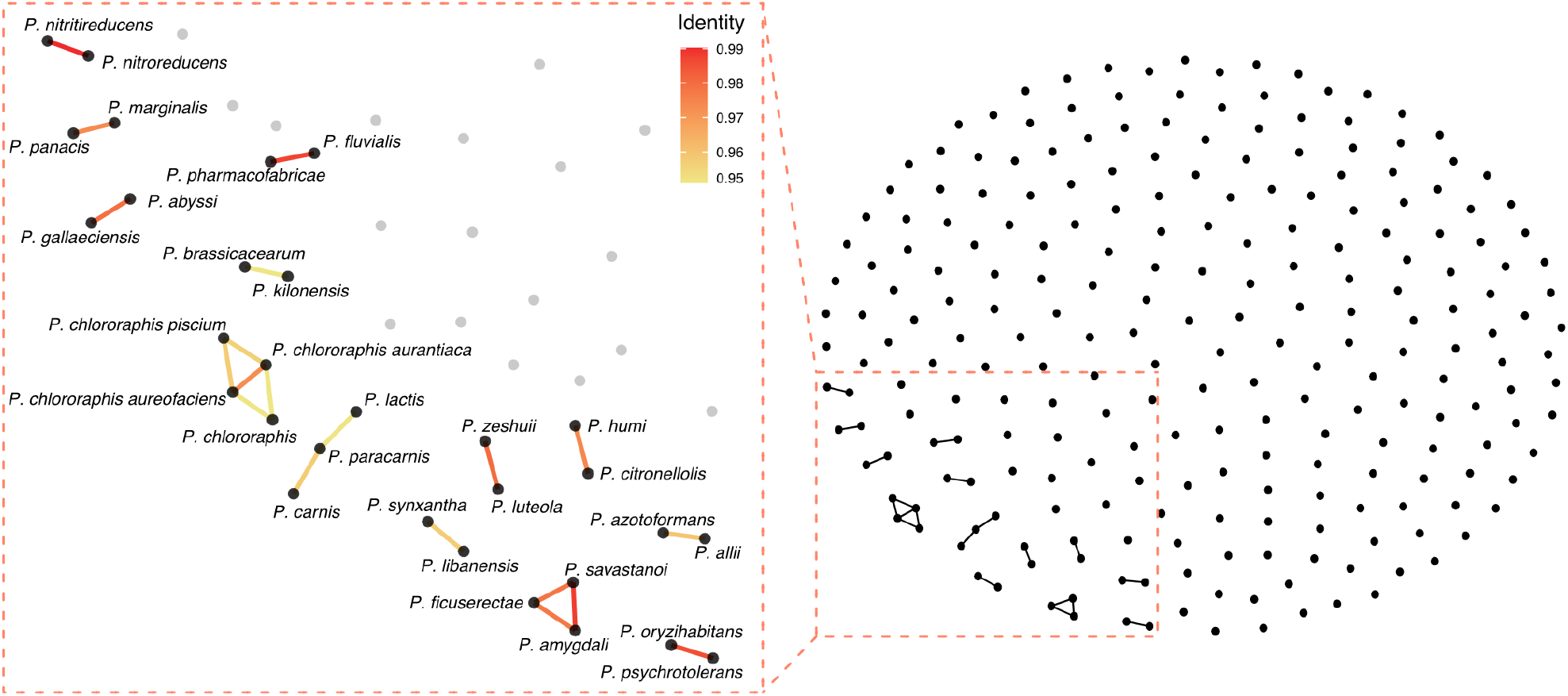
Type strain validation based on Average Nucleotide Identity. Each node in the network represent a type strain genome and nodes are connected if they share at least 95% identity. The left panel is a magnified representation of the connected nodes, with edges colored according to percent identity between the nodes.

We identified 30 connected genomes in the ANI network (Figure 1). Four of these connected genomes are expected because they represent *P. chlororaphis* and its subspecies. Of the 26 remaining connected species, 15 have been previously reported, such as that in the group containing *P. amygdali*, *P. ficuserectae*, and *P. savastanoi*^15, 16^. Here, we observed 11 connections, including the one between *P. panacis* and *P. marginalis* with 97.34% identity, suggesting that *P. panacis* is a later synonym of *P. marginalis*.

### The *Pseudomonas* genomic distance network is highly structured

In networks, the community structure plays an important role in understanding network topology. We used all 10,035 *Pseudomonas* genomes to construct a distance network to estimate the number of *Pseudomonas* species from the number of communities detected in this network. Since alignment-based methods to estimate genome similarity (e.g. ANI) is computationally expensive due to the algorithm quadratic time complexity^19^, it becomes impractical for thousand genomes. Therefore, we estimated the Mash distance that strongly correlates with ANI and can be rapidly computed for large datasets^20^.

Mash distances are computed by reducing large sequences to small and representative sketches^20^. We estimated the pairwise Mash distance for all genomes using sketch sizes of 1000 and 5000, which converged to similar distance values (Figure S2a). However, we observed that the greater the distance between two *Pseudomonas* genomes, the more divergent the distance estimation (Figure S2b), although the density distribution is similar (Figure S2c). The final distance between two genomes was given as the average distance value from both sketch sizes. We used the reciprocal Mash distance (1 - Mash) to estimate the ANI for all 10,035 genomes.

We generated a weighted *Pseudomonas* distance network considering nodes as genomes and edges as the identity between two genomes. Although the 95% ANI value has been widely accepted to delineate species, we evaluated how different thresholds affect network topology by assessing density, transitivity, and the number of connected components (Figure 2). The network density, i.e., the ratio of the number of edges and the number of possible edges, decreased throughout the interval but stabilized between 90% and 97% ANI, keeping the network topology almost unchanged (Figure 2a). To estimate how structured the network was with different ANI thresholds, we also computed the average network transitivity (also called average clustering coefficient) (Figure 2b). The average transitivity is the normalized sum over all local transitivities (the probability of a given node having adjacent nodes interconnected). The high transitivity values revealed that the *Pseudomonas* network is highly structured (i.e., formed by tightly connected clusters) (Figure 2b). This structured profile was observed before for the *P. putida* group network^17^, indicating that communities in *Pseudomonas* distance networks rarely overlap.

**Figure 2.**
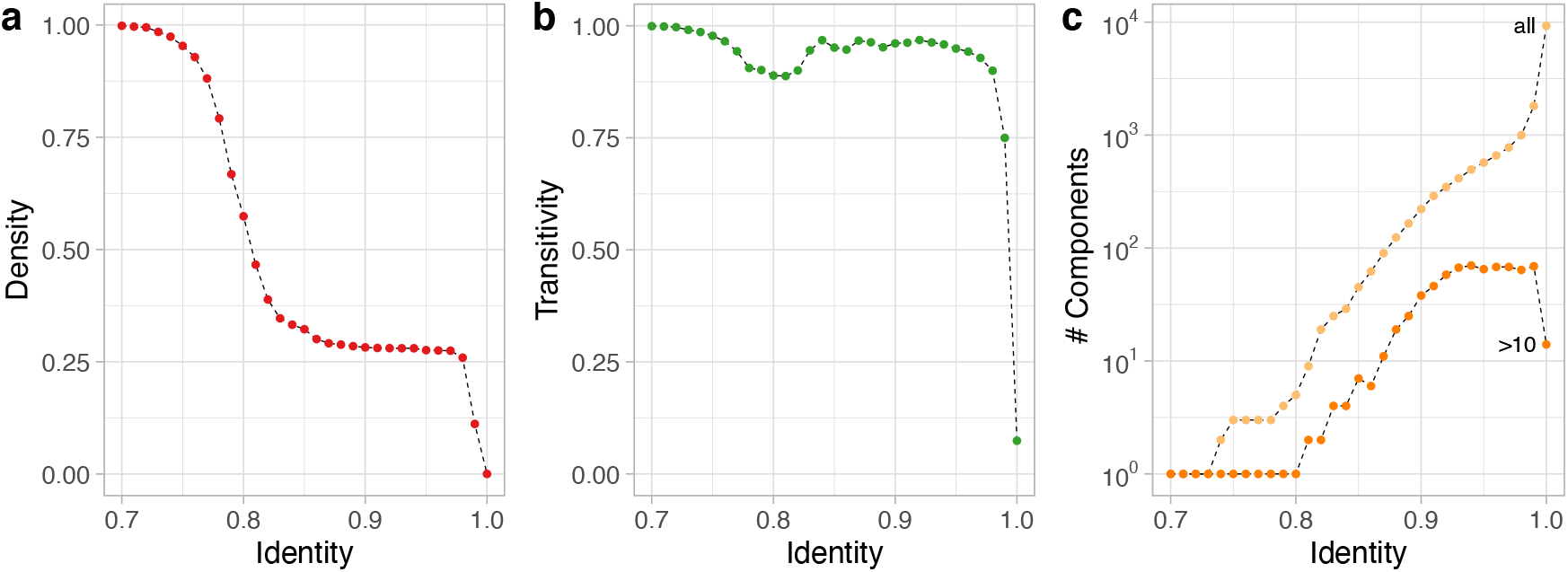
*Pseudomonas* distance network topology evolution. a) Proportion of present connections (network density) and b) average transitivity change over different identity (1 – Mash) cut-off values. c) Number of network components detected with different identity thresholds. Light orange dots represent the total number of components, whereas the dark dots represent only components with more than ten nodes.

To decrease the influence of overrepresented species (e.g., *P. aeruginosa*) on the topological network statistics, we also computed the variation in the number of components (Figure 2c). A connected component in a network is a subset of nodes connected via a path. At 70% identity, we had a single giant connected component. Expectedly, the number of connected components increased with the identity threshold because of the emergence of smaller components or even orphan nodes. Interestingly, connected components with more than ten nodes arose only above 81% identity threshold and stabilized close to 95%, highlighting that the 95% ANI threshold is accurate for species demarcation.

We used the *Pseudomonas* network discarding connections lower than 95% identity to estimate the number of species from the number of communities in the network. We detected 573 communities by using the label propagation algorithm^21^. This number is similar to the number of connected components at 95% identity threshold (n = 570), further supporting that the *Pseudomonas* distance network is highly structured, containing non-overlapping communities. By considering each community as a different *Pseudomonas* species, we evaluated the distribution of type strains in these communities.

Seventeen communities had more than one type strain in the same cluster, indicating the existence of later heterotypic synonyms, as shown in Figure 1. For each community, we assigned only one representative genome (see methods for more detail). For example, in the community containing *P. amygdali, P. ficuserectae, and P. savastanoi*, we maintained *P. amygdali* as the representative strain and the others were considered later heterotypic synonyms, as previously proposed^11^. We observed that only 210 communities (36.64%) had representative genomes from validly described species, reinforcing the underestimation of the number of *Pseudomonas* species if only the type strains are considered.

Regarding the community’s sizes, *P. aeruginosa* corresponds to the largest community, comprising 5116 genomes (Figure 3, Table S3). Most communities had few genomes. Although large communities tend to have type strains, 61 type strains (29.04%) are single nodes (Figure 3, Table S3), further demonstrating that estimating the diversity of *Pseudomonas* only by type strains severely underestimates diversity. For example, the community containing *Pseudomonas spp7* has 122 genomes and is potentially a new genomospecies.

**Figure 3.**
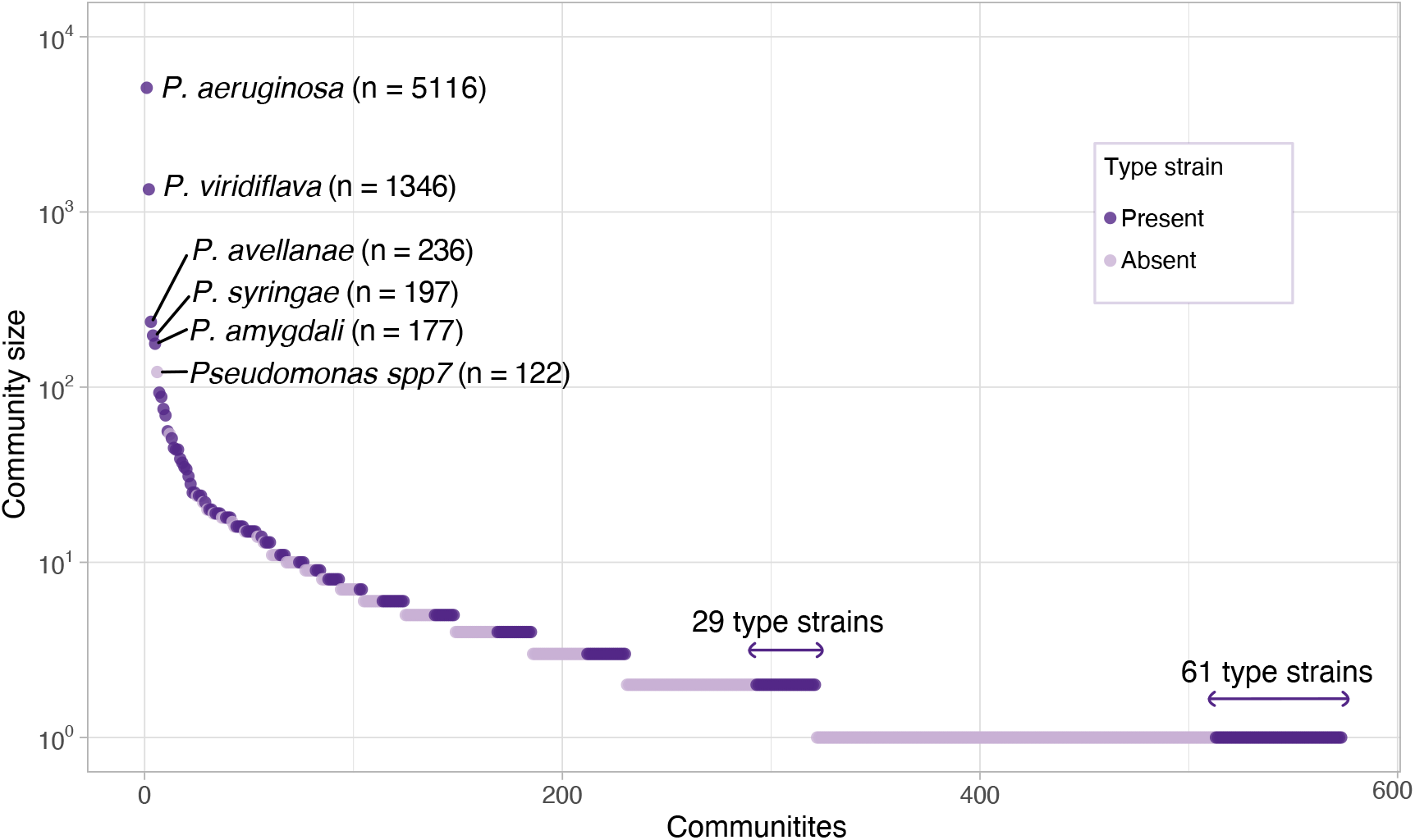
*Pseudomonas* community sizes. Dark and light purple dots represent communities with and without type strains, respectively. The names and number of genomes are displayed in those communities with more than 100 genomes. y-axis is in log scale.

### Comparison with NCBI classification highlights *Pseudomonas* misclassification

After delimiting the species by the community detection approach, we compared them with the classification available in NCBI Taxonomy^22^. Briefly, we computed how many genomes were deposited with a given species name and how many genomes were identified for that species by our network approach. Of the 10,035 genomes used in this work, 25.65% were misclassified in NCBI Taxonomy (Table S5). This proportion includes species considered as later synonyms that should be reclassified (e.g. *P. savastanoi*), non-classified genomes (*Pseudomonas sp.)*, and those genomes that are unconnected to the expected species cluster. The most poorly classified species were *P. brassicacearum* (95.65%), *P. fluorescens* (95.23%), *P. stutzeri* (94.58%), and *P. putida* (88.70%). This high rate of misclassification is linked to the type strain determined for each species. For example, the critical classification problem of *P. putida* has been recently reported by us^17^. The *P. putida* NBRC 14164^T^ type strain forms an isolated community in the network with only 15 genomes. On the other hand, the community of *P. alloputida* Kh7^T^ harbors 69 genomes, constituting the largest community in the *P. putida* group. Thus, most of the genomes deposited as *P. putida* are actually from *P. alloputida*. Regarding the misclassification of *P. stutzeri*, 122 genomes fall into the community represented by *Pseudomonas spp7*, a potentially new genomospecies mentioned above.

We also assessed the impact of our approach defining the species-level taxonomy of the 1,083 non-classified *Pseudomonas* genomes available in Genbank (*Pseudomonas sp*.). Interestingly, 511 *Pseudomonas sp.* genomes (47.18%) were distributed among 97 communities containing type strains (Table S6). The species that received the most genomes were *P. glycinae* (n = 35), *P. lactis* (n = 34), and *P. mandelii* (n = 31).

### The *Pseudomonas* phylogeny reveals at least fourteen groups

To reduce the influence of overrepresented species, we used the 573 representative genomes from each community to retrieve orthologous genes and reconstruct the *Pseudomonas* phylogeny. The *Cellvibrio japonicus* Ueda 107^T^ was used as an outgroup. We identified 31,094 orthogroups, of which 168 were present in all species, including 30 single-copy genes. We used the single-copy genes to reconstruct the *Pseudomonas* phylogeny and identify the main *Pseudomonas* groups (Figure 4).

**Figure 4.**
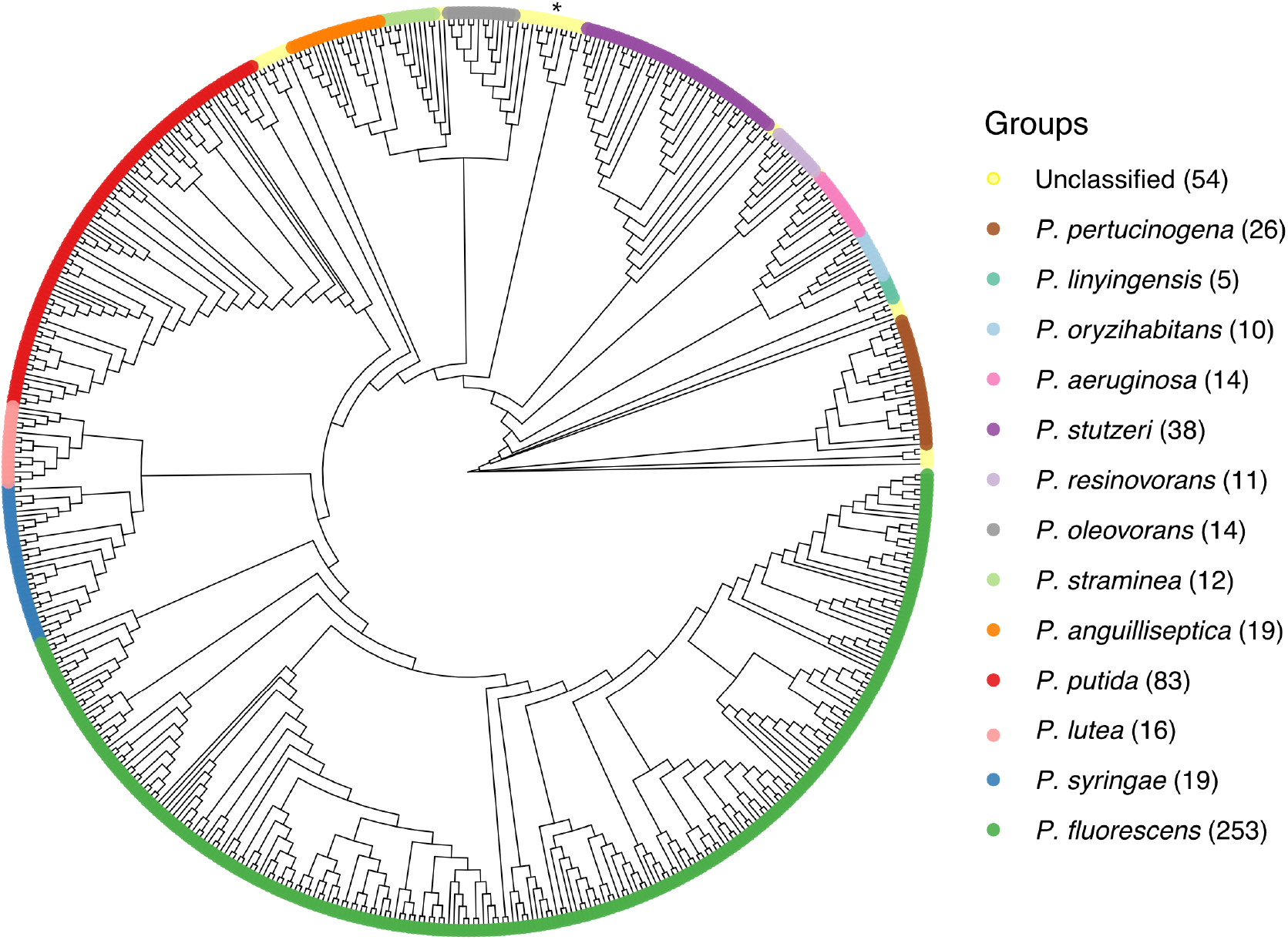
Phylogenetic tree mapping *Pseudomonas* groups. Maximum-likelihood phylogenetic tree using core single-copy genes in representative genomes from 573 communities detected in the *Pseudomonas* network. Colors indicate *Pseudomonas* groups. The number of genomes in each group is in parenthesis. The asterisk highlights the *P. alcaligenes* group described here. The outgroup is *Cellvibrio japonicus* Ueda 107^T^.

The main *Pseudomonas* groups have been previously characterized using housekeeping genes such as 16S rDNA, *gyrB*, *rpoB*, and *rpoD* from type strains ^8, 15^. To delineate each group, we retrieved those representative genomes (species) within previously-described groups (Table S7). We then tracked the Most Recent Common Ancestor (MRCA) for those species in the *Pseudomonas* phylogenetic tree to include uncharacterized representative genomes as well. For example, the *P. lutea* group comprises three known species: *P. abietaniphila*, *P. graminis*, and *P. lutea*^8^. By tracking the corresponding MRCA node, we ensured the monophyly and included *P. bohemica* and 12 uncharacterized species in this group (Table S3). This approach allowed a more accurate characterization of both recently described type strains and other uncharacterized species (Figure 4, Table S3). We identified the 13 main *Pseudomonas* groups and one new group with 10 genomes and three type strains: *P. alcaligenes*, *P. fluvialis*, and *P. pohangensis* (Figure 4, Table S8). Since *P. alcaligenes* is the firstly-described type strain in this group^23^, we named this group as *P. alcaligenes* group.

### Lineage and genus boundaries

The genus *Pseudomonas* has three recognized lineages: *P. pertucinogena*, *P. aeruginosa*, and *P. fluorescens*. The *P. pertucinogena* lineage is composed of a single phylogenetic group. The *P. aeruginosa* lineage comprises 6 phylogenetic groups (*P. oryzihabitans*, *P. stutzeri*, *P. oleovorans*, *P. resinovorans*, *P. aeruginosa*, and *P. linyingensis*). The *P. fluorescens* lineage also comprises 6 phylogenetic groups (*P. fluorescens*, *P. lutea*, *P. syringae*, *P. putida*, *P. anguilliseptica*, and *P. straminea*); the *P. fluorescens* group is further divided into 8 or 9 phylogenetic subgroups^15^. In this work, 70.38% of the communities (species) belong to the *P. fluorescens* lineage, 16.72% to *P. aeruginosa*, and 4.52% to *P. pertucinogena*; 8.36% were unclassified communities. We observed that, unlike the *P. pertucinogena* and *P. fluorescens* lineages, the *P. aeruginosa* lineage is polyphyletic (Figure 5a).

**Figure 5.**
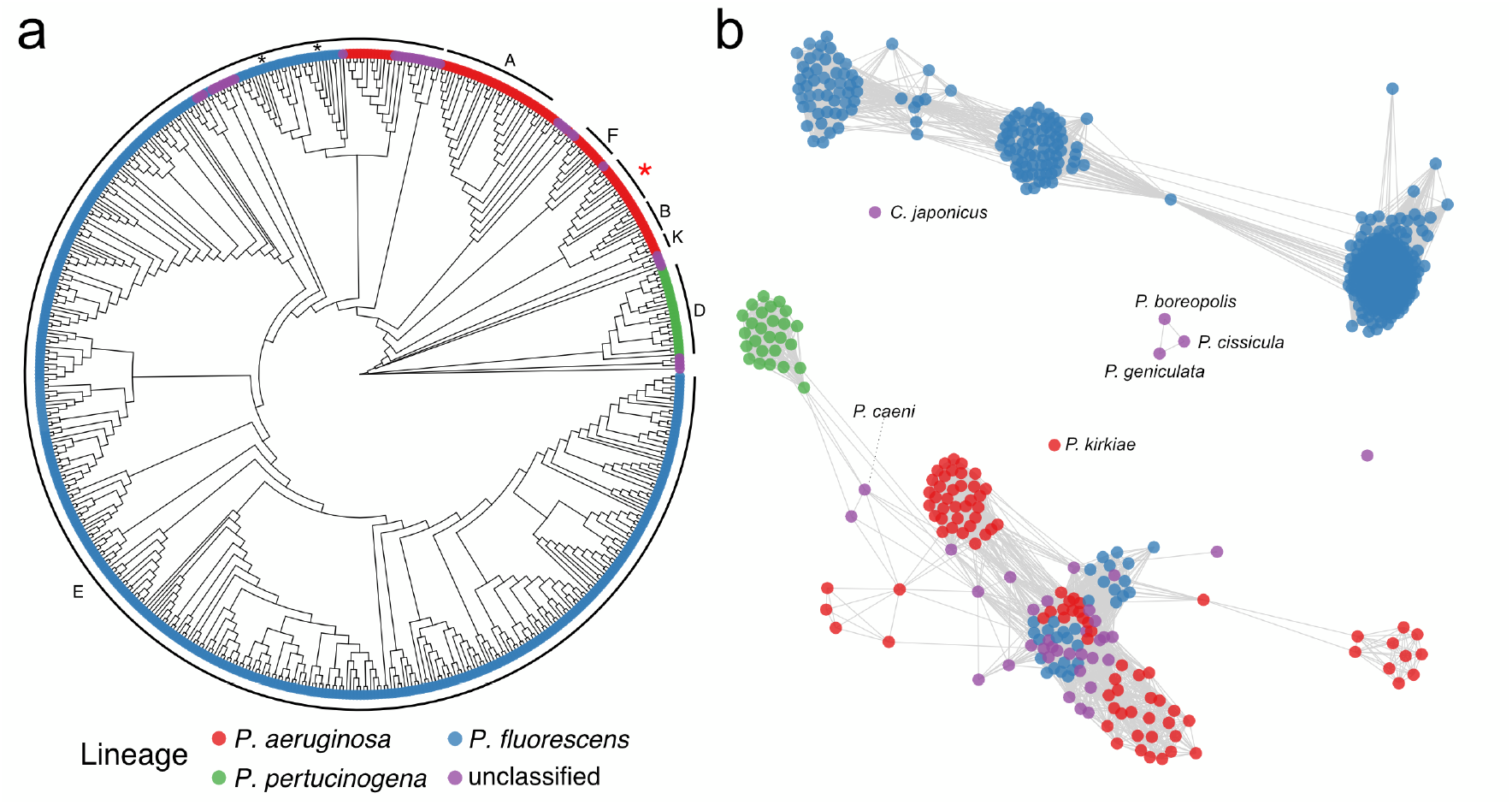
*Pseudomonas* phylogenetic tree with proposed genus boundaries and Percentage of Conserved Proteins (POCP) network. a) Phylogenetic tree annotated with *Pseudomonas* lineages. The outer letters indicate the annotation adopted by the Genome Taxonomy Database (GTDB). The genus proposed to keep the name *Pseudomonas* is marked with a red asterisk. Other genera proposed by GTDB adopt the nomenclature “*Pseudomonas*” followed by a letter (e.g. *Pseudomonas_E*); for clarity, only the letters and those proposed genera with more than five communities are displayed. b) Network based on POCP index using a 60% threshold. Colors represent lineages. Blue nodes embedded in the component with genomes of *Pseudomonas aeruginosa* lineage belong to the groups *P. anguilliseptica* and *P. straminea*; these two groups are marked in the phylogenetic tree with black asterisks.

We used the Genome Taxonomy Database (GTDB) approach^13^ to evaluate whether *Pseudomonas* should be divided into different genera. The GTDB proposes a framework to classify genomes in higher taxonomic ranks (e.g. genus). By using the GTDB classification, *Pseudomonas* should be divided into 17 genera named generically with “Pseudomonas” followed by a letter (e.g. “*Pseudomonas_A*”), with the *P. aeruginosa* group retaining the name *Pseudomonas*. We found a high correspondence between *Pseudomonas* groups and the proposed genera, with few inconsistencies (Figure 5a, Table S8). According to the GTDB classification, the *P. fluorescens* lineage, together with the *P. oleovorans* group and the here described *P. alcaligenes* group, would form a single genus called *Pseudomonas_E* (Figure 5a), which corresponds to 77.52% of the species (communities) estimated in our study.

We also used the Percentage of Conserved Proteins (POCP) index to evaluate the relationships between lineages (Figure 5b) and complement the GTDB approach. Briefly, the POCP index measures the proportion of shared proteins between two genomes^24^. The original proposal is that genomes belong to the same genus if they share at least half of their proteins^24^. By using 50% as a threshold, we observed that only the outgroup *C. japonicus* and other four genomes do not belong to the main POCP network component with all lineages. However, we observed two main clusters by using a 60% threshold to link communities (Figure 5b).

Apart from *P. anguilliseptica* and *P. straminea* groups, the *P. fluorescens* lineage forms an isolated component in the network (Figure 5b). The *P. pertucinogena* and *P. aeruginosa* lineages are in the same component, but linked by a few connections, including a bridge via a *P. caeni* genome. The outgroup *C. japonicus* is an orphan in the network, as well as *P. kirkiae*. The species *P. boreopolis*, *P. cissicula*, and *P. geniculata* were also isolated. These three species have already been recognized as belonging to the genus *Xanthomonas*^25^. Nevertheless, they remain classified as *Pseudomonas* in Genbank and are still labeled as validly published with a correct name in LPSN.

## DISCUSSION

The *Pseudomonas* genus underwent several taxonomic reclassifications over the years. Here, we used 10,035 *Pseudomonas* genomes to estimate the genus diversity through network analysis and community detection. We observed that several type strains are later synonyms and should be officially revised, as also noted elsewhere^8, 15^.

Regarding the *Pseudomonas* network, we observed that the number of detected communities is very close to the number of network components at a 95% identity threshold. Combined with the stabilization of density and high transitivity around this threshold, we conclude that the *Pseudomonas* network is highly structured. This structured network profile has also been noted previously reported for the *P. putida* group^17^.

Considering each community as a different genomospecies, we identified 573 communities, way more than the 233 *Pseudomonas* species with validly published names. Moreover, we found 61 orphan type strains in the network, indicating that the diversity estimated using only type strains is highly underestimated. In addition, this work shows that 25.65% of the *Pseudomonas* genomes are misclassified. This is a matter of concern, as misclassified genomes in public repositories can introduce noise to pangenome studies, reduce strain typing accuracy, and propagate labeling errors to several studies, including those reporting the characterization of new species.

Here, we also showed potential new genomospecies. For example, the community assigned as *Pseudomonas spp7* contains 122 genomes, and it is a sister group of *P. stutzeri*. The high misclassification rate of *P. stutzeri* (Table S5) can be explained by the presence of this new closely related species. Such inconsistencies could be mitigated through a standardized taxonomic framework, as previously proposed^18^. However, there is still resistance to define species based solely on genome sequences, even with the massive number of available genomes ^18^. Therefore, isolating and characterizing members from *Pseudomonas spp7* community will allow the consolidation of this new species.

Although previous works provided insights about what would be considered *Pseudomonas*^8, 15, 26^, how to delimit the *Pseudomonas* genus remains an open question. We tried to address this problem by using GTDB classification and POCP index network, two approaches proposed to delimit genera. The GTDB results indicate that the *P. fluorescens* lineage and the *P. oleovorans* and *P. alcaligenes* groups would constitute a genus with the generic name *Pseudomonas_E* (Figure 4). However, the POCP index network at 60% shows that *P. straminea* and *P. anguilliseptica* groups are closer to *P. aeruginosa* than to *P. fluorescens* lineage (Figure 4b). Aiming for a parsimonious separation, we propose that the *P. fluorescens* lineage, excluding the *P. straminea* and *P. anguilliseptica* groups, should be considered a new genus. Furthermore, by the GTDB results, the *Pseudomonas* groups from *P. aeruginosa* lineage should also be revised to assess whether they are new genera, as the *P. aeruginosa* lineage itself is polyphyletic. Prioritizing the GTDB approach here should provide the best approach because it normalizes taxonomic ranks and ensures group monophyly^13^.

## CONCLUSION

In this study, we estimated the *Pseudomonas* diversity using a network approach. We show that type strains represent less than half of the estimated number of species, and that many of them are orphans in the network. We discovered new genomospecies and groups, such as *Pseudomonas spp7* and *P. alcaligenes*, respectively. Although genus delineation is somewhat complex, we propose the *Pseudomonas* genus division by combining GTDB classification and POCP index. To fully understand the *Pseudomonas* diversity, it will be important to focus on each group and characterize species from communities without type strains. This study provides a state-of-the-art classification to delimit bacterial species, which we expect to serve as a guide for future studies with *Pseudomonas spp*, reducing the problems caused by misclassified genomes.

## METHODS

### Dataset collection and annotation

We recovered 11,025 genomes of *Pseudomonas* from Genbank in June 2020. Genome quality was evaluated with BUSCO v4.0.6^27^ using the *Pseudomonadales* dataset. We defined completeness as 100% minus the percentage of missing genes, and contamination as the fraction of duplicated genes. Quality was defined as completeness – 5 x contamination^13^. Genomes with more than 400 contigs were removed, and contigs shorter than 500bp were removed from the remaining genomes. We used mash v2.2.2^20^ to calculate the pairwise distances between those genomes with quality higher than 80% using sketches of 1000 and 5000. Regarding the type strains, we used all species with available genomes and validated taxonomic names according to the LPSN^10^ on March 2021. The pairwise distances between type strains were performed using pyani v0.2.10^28^. We reannotated the genomes with prokka v1.14^29^ to allow a systematic large-scale genome comparison.

### Network analysis

By using the pairwise Mash distances, we generated the corresponding graph and obtained the topological graph properties such as density, transitivity, and number of components with the igraph package^30^. We used the label propagation algorithm to detect communities^21^. The representative genome for each community was defined based on three conditions: i) if the community has only one type strain, the type strain was considered the representative genome; ii) if the community has more than one type strains, the first described type strain was chosen; iii) else, we randomly chose a genome in a community (seed = 1996) and assigned the community name with the notation *Pseudomonas sppX*, where *X* is the community number.

### Phylogeny and POCP index

We used OrthoFinder v2.5.2^31^ to obtain the orthogroups from community type genomes. All single-copy genes were aligned with MAFFT v7.467^32^ and concatenated to reconstruct the *Pseudomonas* phylogeny with IQ-TREE v2.1.2^33^. The best-fit model detected through ModelFinder ^34^ was LG+F+I+G4. One thousand bootstrap replicates were generated to assess the significance of internal nodes. Phylogenetic trees were visualized and annotated using ggtree^35^. We tracked MRCA nodes for *Pseudomonas* groups definitions using treeio^35^.

The Percentage of Conserved Proteins (POCP) between two genomes were calculated using the formula 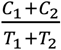, where *C* is the number of conserved proteins and *T* is the total number of proteins^24^. The number of conserved proteins was obtained from the orthologs matrix *A*_ij_ generated by OrthoFinder, where each entry (*i*, *j*) is the total number of genes in species *i* that have orthologues in species *j*. The graphs were generated and visualized using igraph^30^ and ggnetwork v0.5.8^36^, respectively. The GTDB classification was obtained in April 2021 (http://gtdb.ecogenomic.org/).

## ACKNOWLEDGEMENTS

This work was supported by Fundação Carlos Chagas Filho de Amparo à Pesquisa do Estado do Rio de Janeiro (FAPERJ; grants E-26/203.309/2016 and E-26/203.014/2018), Coordenação de Aperfeiçoamento de Pessoal de Nível Superior - Brasil (CAPES; Finance Code 001), and Conselho Nacional de Desenvolvimento Científico e Tecnológico. The funding agencies had no role in the design of the study and collection, analysis, and interpretation of data and in writing.

## SUPPLEMENTARY FIGURES

**Figure S1.**
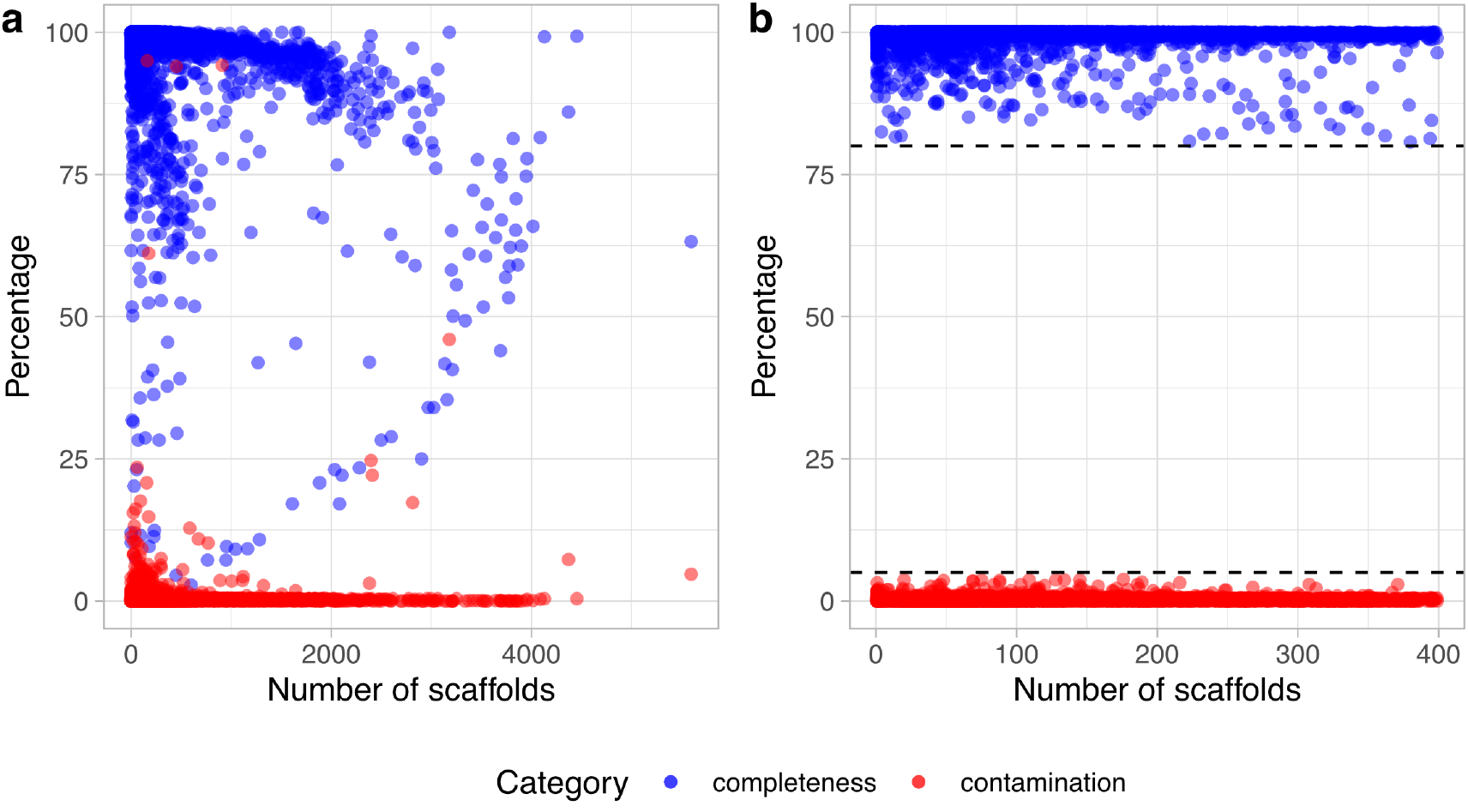
BUSCO estimation for completeness and contamination for all *Pseudomonas* genomes. a) Distribution for all 11,025 *Pseudomonas* genomes. b) Genomes used in this study after discarding genomes based on 80% quality threshold and fragmentation higher than 400 scaffolds (see methods).

**Figure S2.**
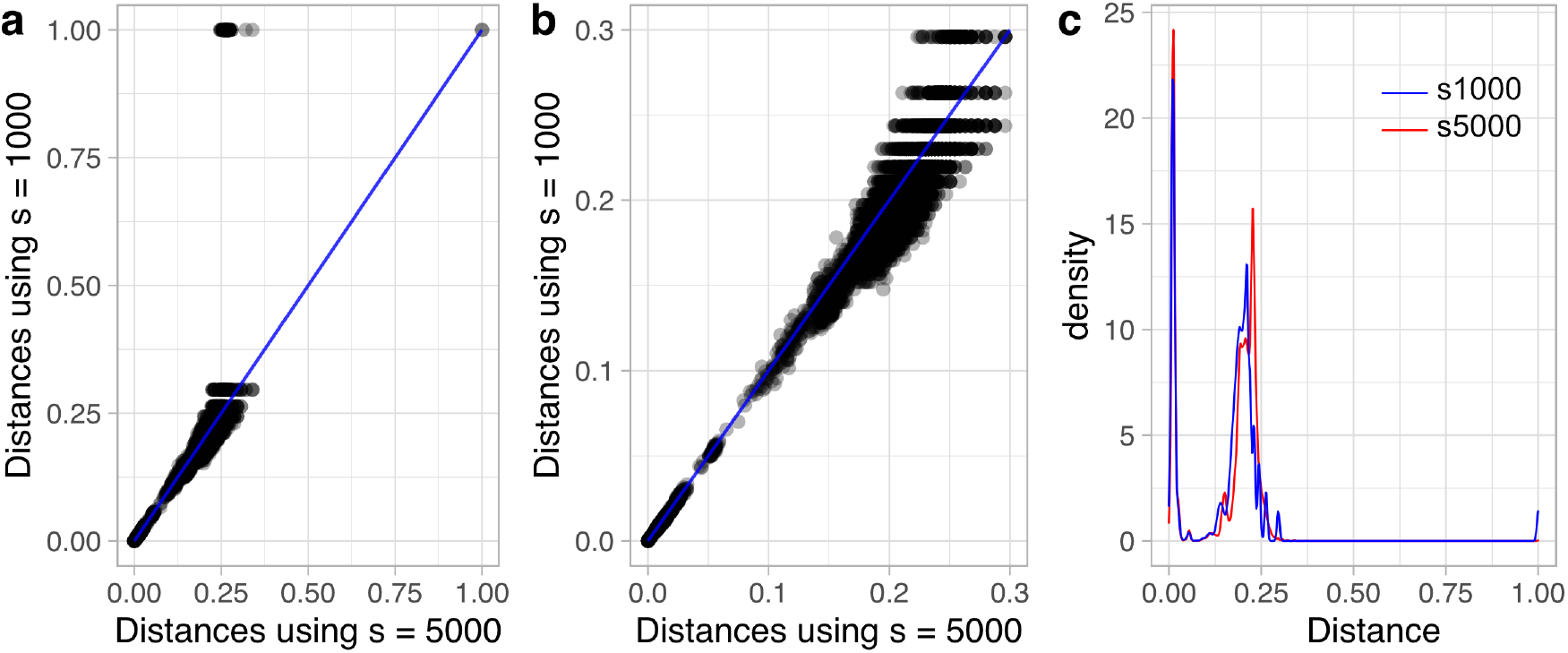
Mash distance statistics. a) Comparison of estimated Mash distance using sketches sizes of 1000 and 5000. b) Mash distances restricted to the interval [0.0, 0.3] in both axes. c) Mash distance distribution for each sketch size.

**Figure S3.**
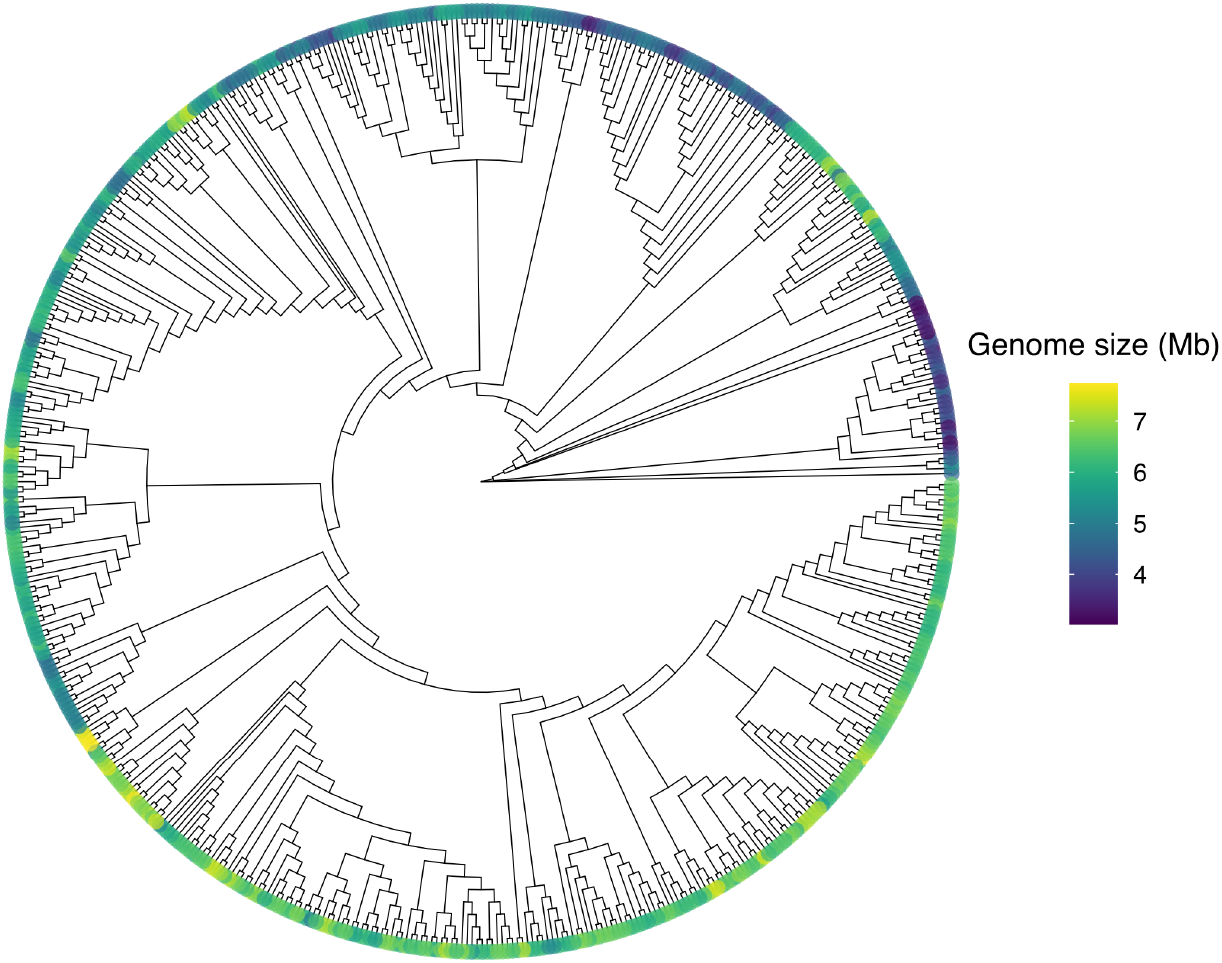
Genome size distribution for *Pseudomonas* communities. Maximum-likelihood phylogenetic tree using core single-copy genes in representative genomes from 573 communities detected in the *Pseudomonas* distance network. Colors indicate the genome size distribution.

